# Microbial community profiles of the snake cloaca in the presence and absence of *Chlamydiota*

**DOI:** 10.64898/2026.06.04.730152

**Authors:** Ehsan Ghasemian, Sedreh Nassirnia, Trestan Pillonel, Sebastien Aeby, Simon Rüegg, Claire Bertelli, Nicole Borel, Gilbert Greub

## Abstract

*Chlamydiota* are obligate intracellular bacteria detected in snake cloacal microbiota, yet their biological significance remains poorly understood. Members range from recognised pathogens, such as *Chlamydia serpentis*, to potential environmental symbionts, raising questions about whether they represent transient contaminants, persistent colonisers, or subclinical infectious agents. Despite the cloaca serving as a primary site of chlamydial shedding in snakes, its interaction with the broader cloacal microbiota remains unexplored. Following pan-*Chlamydiota* PCR screening of 137 captive snakes across five collections, 52 samples (caenophidian snakes) (27 *Chlamydiota*-positive, 25 *Chlamydiota*-negative) were retained after V3-V4 16S rRNA sequencing and quality filtering. Presence of *Chlamydiota* was not associated with significant differences in alpha diversity or overall community composition, though it was related to greater within-community compositional heterogeneity. Differential abundance and multivariate analyses identified several enriched and depleted genera, with *Lachnospiraceae* and *Copromonas* consistently negatively associated with *Chlamydiota* across all three methods. Co-occurrence network analysis recovered more associations and a higher proportion of positive edges in the presence of *Chlamydiota*, with an expansion of anaerobic taxa. Inferred functional composition did not differ globally between groups; however, elastic net stability selection identified subtle pathway-specific differences, including enrichment of proteolytic and mycobacterial pathways in infected snakes. Our findings suggest subtle infection-associated community shifts that do not fully conform to established mammalian paradigms in which *Chlamydia* species behave either as gastrointestinal commensals or as cervicovaginal pathogens, highlighting the need for multi-omics approaches in larger cohorts of caenophidian and henophidian wild and captive snakes to better characterise the mechanistic basis and generalisability of these associations.

## Introduction

*Chlamydiota* are obligate intracellular Gram-negative bacteria characterised by a unique biphasic lifecycle alternating between the infectious elementary body (EB) and the replicating reticulate body (RB) [1,2]. Whilst historically associated with human and mammalian diseases [3,4], accumulating evidence over the past two decades has demonstrated that snakes harbour chlamydial species [5–10]. Early studies reported *Chlamydia pneumoniae*, originally described as a human respiratory pathogen, as the predominant chlamydial agent in both free-ranging and captive reptiles, causing granulomatous lesions in visceral organs [8]. Subsequent molecular and culture-based investigations have since revealed a far broader chlamydial diversity in snakes [5,7,9]. Through metagenomic sequencing, Taylor-Brown *et al*. [6] characterised *Candidatus* Chlamydia sanzinia from the choana of a captive Madagascar tree boa. Staub *et al*. [7] described two additional novel species, *Chlamydia serpentis* and *Chlamydia poikilothermis*, isolated from cloacal and choanal swabs of captive snakes from Swiss collections, demonstrating that isolation success and growth characteristics were temperature-dependent. *C. serpentis* has since emerged as a clinically relevant pathogen, with outbreak reports in zoological collections documenting sudden deaths, granulomatous inflammation, and fibrinous pneumonia [5]. Longitudinal surveillance further highlighted the complex epidemiology of *C. serpentis*, with evidence of persistent infection, intermittent shedding, and environmental persistence contributing to recurrent outbreaks within the same collections [11]. Cloacal samples have been central to the detection and monitoring of chlamydial infections across these studies [5,7,8], suggesting the cloaca as a primary site of chlamydial shedding in snakes.

*Chlamydia* has long been recognised as an inhabitant of the gastrointestinal (GI) tract across a remarkably diverse range of host species. In cattle, sheep, goats, pigs, and birds, *Chlamydia* colonise the lower intestinal tract, particularly the caecum and colon, for months to years in the complete absence of notable pathological changes, exhibiting a commensal relationship characterised by continuous faecal shedding and faecal-oral transmission [12]. In the murine model, oral or intravaginal inoculation with *Chlamydia muridarum* establishes GI colonisation lasting 100-285 days, whereas infection at other mucosal sites resolves within weeks; this persistence may be attributed to immune downregulatory mechanisms in the gut that suppress the local adaptive response and prevent inflammatory cell recruitment, allowing the pathogen to persist beneath the protective mucin layers of the intestinal epithelium [12,13]. At the cellular level, long-term GI colonisation in mice disrupts intestinal homeostasis by impairing goblet cell differentiation, suppressing tight junction integrity, and dysregulating immune-epithelial crosstalk, mechanisms that may facilitate pathogen persistence and barrier dysfunction [14]. In the female reproductive tract, the vaginal microbiome profoundly influences susceptibility to *C. trachomatis* infection: whilst *Lactobacillus crispatus*-dominated communities confer protection through lactic acid-mediated chlamydial killing, dysbiotic communities enriched in anaerobes and indole-producing taxa such as *Prevotella* species (spp.) facilitate immune evasion by supplying the indole precursor required for tryptophan synthase-dependent rescue of *C. trachomatis* from interferon (IFN)-𝛾-induced persistence [15–17].

Snake cloaca serves not only as a site of faecal and urinary excretion but also as a convergence point for reproductive, digestive, and urinary tracts [18]. The cloacal microbiota of snakes comprises a diverse community shaped by host species, diet, phylogeny, and environmental conditions dominated at the genus level by *Bacteroides*, *Cetobacterium*, *Salmonella*, *Fusobacterium*, and *Clostridium* [19–24]. Of note, the phylum *Chlamydiota* has been detected in the gut and oral microbiota of multiple snake species across independent metagenomic studies. In a survey of four snake species from southern China, *Chlamydiota* was detectable at the phylum level within the gut microbiota, with their relative abundance varying by host species [21]. Subsequently, Hu *et al*. [20] reported *Chlamydiota* exceeding 0.68% relative abundance in the oral samples of three Chinese snake species examined, whilst members of this phylum also appeared in faecal samples. In addition, Zhu *et al*. [25] identified *Simkaniaceae* (family within the phylum *Chlamydiota*) as a dominant bacterial group, comprising over 10% relative abundance, in the gut microbiota of the invertebrate-feeding *Rhabdophis pentasupralabralis*, representing the first report of *Simkaniaceae* dominance in snake microbiota studies. Song *et al*. [26] similarly found *Chlamydiota* abundance to be significantly elevated in *Rhabdophis chiwen* compared to other snake species, with this enrichment potentially linked to dietary exposure through soil invertebrates.

The biological significance of *Chlamydiota* within snake cloacal microbiota remains unclear; members of this phylum encompass both obligate intracellular pathogens and environmental symbionts, and their detection in metagenomics datasets raises questions regarding whether they represent transient environmental contaminants acquired through prey, persistent cloacal symbionts, or subclinical infectious agents capable of modulating host immune and microbial community dynamics. Given that the cloaca is a site of intermittent bacterial shedding, understanding how *Chlamydiota* interact with the broader cloacal microbial community represents a yet underexplored dimension of snake microbial ecology. To date, most studies addressing host microbiota-*Chlamydiota* interactions have focused on the human cervicovaginal microbiota in the context of *C. trachomatis* infection [27–31], with only a limited number of investigations in other mammalian systems [32–34]. The snake, as a phylogenetically distant reptilian host with a distinct anatomical and physiological cloacal niche, therefore offers an underexplored model from which new aspects of this interaction may be uncovered. This study was conducted within the framework of a *Chlamydiota* screening investigation in snakes, in which molecular diagnosis of *Chlamydiota* was the primary objective [9]. Nevertheless, this prospective cohort allowed us to explore the cloacal microbiota of caenophidian snakes in the presence and absence of *Chlamydiota*.

## Materials and Methods

### Sample collection

Five private collections (anonymised as collections 1–5) contributed in providing cloacal and choanal swabs from 137 snakes. DNA was extracted from all samples according to Borel *et al*. [10]. Of the 137 snakes initially screened by pan-*Chlamydiota* real-time PCR [35], samples with a cycle threshold (Ct) value ≤ 35 were classified as *Chlamydiota*-positive, those with no detectable amplification (no recorded Ct value) were classified as *Chlamydiota*-negative, and samples with a Ct value > 35 were excluded as inconclusive. Applying these criteria yielded 99 snakes (59 *Chlamydiota*-positive and 40 *Chlamydiota*-negative), which were subsequently selected for 16S rRNA amplicon sequencing and microbiota profiling.

16S rRNA sequencing Library preparation followed the Illumina 16S Metagenomic Sequencing Library Preparation protocol [36]. Briefly, the V3-V4 hypervariable region of the bacterial 16S rRNA gene was amplified by a two-step PCR: an initial 25-cycle amplification followed by an 8-cycle indexing PCR to incorporate barcodes and Illumina adapters. Libraries were normalised, pooled, and sequenced on an Illumina NextSeq 1000 (P1 v3 kit; 2 x 150 bp paired-end reads) with 10% PhiX spike-in. We included a commercial mock community (MSA-2002^TM^, ATCC, Manassas, USA) as a positive control to assess sequencing success, an extraction blank as a negative control (NEC) to detect reagent-level contamination introduced during DNA extraction, and negative library controls (NLCs) to identify contamination arising during library preparation.

### Data processing

Raw 16S rRNA gene sequencing reads were processed using the in-house zAMP pipeline (v1.0.0) [37], which integrates quality assessment (FastQC), adapter trimming and read filtering (Cutadapt), error correction and amplicon sequence variant (ASV) inference (DADA2), and taxonomic classification against the Greengenes2 database (v2024.09). Sequencing generated 4,329,385 raw reads across the samples, of which 3,314,637 were retained after quality filtering and 3,238,912 reads remained following chimera removal (Table S1).

Putative contaminant taxa were identified with the decontam package (v1.16.0) using the prevalence-based approach (threshold = 0.1), which contrasts ASV prevalence in NEC/NLCs against that in true samples. To avoid the inadvertent removal of biologically meaningful taxa, a rescue criterion was implemented whereby ASVs initially flagged as contaminants were retained when their mean relative abundance was greater in true samples than in NEC/NLCs (ratio ≥ 2). After decontamination, taxa and samples with zero total counts were pruned from the phyloseq object.

To account for uneven sequencing depth across samples, a consensus rarefaction approach was employed. The rarefaction depth was chosen by inspecting sample-wise rarefaction curves for observed ASV richness, Chao1, Shannon and Simpson indices generated across a range of subsampling depths. A threshold of 5,000 reads per sample was selected as it represented the point at which all four diversity metrics showed substantially reduced slope, indicating adequate coverage of the underlying community while minimising sample loss; depths beyond this point yielded only marginal gains in observed diversity but resulted in the exclusion of additional samples. All samples were therefore rarefied to a uniform depth of 5,000 reads per sample, and samples (n = 29) falling below this threshold were excluded from downstream analyses. To minimise the stochastic effects inherent in single rarefaction events, we performed 100 independent rarefaction iterations, each using a unique random seed. Consensus abundances were calculated by averaging taxon counts across all 100 rarefaction iterations, with final values rounded to integers.

### Alpha Diversity Analysis

Prior to fitting the alpha diversity models, we evaluated whether sex and husbandry facility were better treated as fixed or random effects. Sex was retained as a fixed effect, because factors with fewer than five levels yield unstable random-intercept variance estimates [38,39]. For husbandry facility (k = 4), the decision was supported by a diagnostic protocol comprising intraclass correlation coefficients (ICC) from null mixed models, boundary-corrected likelihood ratio tests using a 50:50 𝜒^2^ mixture of 0 and 1 degrees of freedom [40], singular-fit checks, and sensitivity analyses contrasting fixed-only and random-intercept specifications. Three findings supported a fixed-effect specification. First, Chao1 returned a low but non-negligible ICC (0.073) and a borderline likelihood ratio test (p = 0.092), and a sensitivity analysis comparing the two parameterisations gave concordant *Chlamydiota*-effect estimates (fixed-effect p = 0.917; mixed-model p = 0.634). Second, Shannon diversity and Pielou’s evenness both yielded singular fits (ICC = 0), indicating negligible facility-level clustering. Third, the small number of facilities (k = 4) lies below the recommended threshold for stable random-effect estimation. Together, these results supported modelling both sex and husbandry facility as fixed effects throughout the alpha diversity analyses.

To assess differences in alpha diversity at ASV level between *Chlamydiota* presence and absence, three metrics were quantified: richness (Chao1), diversity (Shannon index), and evenness (Pielou’s evenness). Each metric was calculated within each of the 100 rarefaction rounds, that is, on the per-round rarefied count tables, prior to any averaging or rounding of counts, and the per-sample mean across rounds was retained for downstream analyses. This ordering ensured that the rare-taxon information required by richness-sensitive estimators was preserved at the point of calculation, rather than being attenuated by averaging and integer rounding of the count tables. Analysis of covariance (ANCOVA) models were then fitted to the per-sample mean values with *Chlamydiota* status as the main explanatory variable. Models were adjusted for potential confounders, including sex and husbandry facility, both specified as fixed effects. Type III sums of squares were used to evaluate the effect of *Chlamydiota* status while accounting for covariates, and estimated marginal means were derived to obtain confounder-adjusted group comparisons. Statistical significance was determined at α = 0.05, and pairwise comparisons were conducted between *Chlamydiota* presence and absence for each diversity metric.

### Beta diversity Analysis

Before performing between-sample comparisons, we determined whether sex and husbandry facility should enter the models as fixed terms or as random intercepts. Sex was treated as a fixed factor, since two categories fall well below the empirical threshold required for reliable estimation of variance components [38,39]. The same conclusion was reached for husbandry facility (k = 4), on three separate grounds. First, design: four facilities lie at or below the lower bound conventionally regarded as sufficient for stable random-effect or stratified-permutation inference. Second, confounding: *Chlamydiota* status and husbandry facility were unevenly distributed across the four facilities (Fisher’s exact test, p = 0.015), and restricting permutations to within-facility strata would have left too sparse a permutation space to evaluate the *Chlamydiota* effect with adequate power. Third, model fit: likelihood ratio tests indicated that adding a facility random intercept did not significantly improve fit relative to the fixed-effect model.

Following ASV-level prevalence (≥10%) and abundance (≥0.01% relative abundance) filtering, counts were collapsed to genus rank, in line with the resolution used throughout the downstream analyses. Pairwise Bray-Curtis and Jaccard distances were derived from this genus-level matrix using phyloseq (v1.54.2) [41]. Compositional differences between *Chlamydiota* presence and absence were then tested by permutational multivariate analysis of variance (PERMANOVA), implemented through the adonis2 function of the vegan package (v2.7.3) [42] with 999 permutations and *Chlamydiota* status, sex, and husbandry facility entered as explanatory terms. Marginal Type II sums of squares (by = "margin") were specified so that each predictor was evaluated conditional on the remaining terms, and the magnitude of each effect was reported as R^2^ together with partial omega-squared (𝜔^2^) to provide a bias-corrected complement. Heterogeneity of multivariate dispersions was subsequently examined with betadisper, followed by a 999-permutation ANOVA and Tukey’s HSD post-hoc contrasts, which together allow genuine shifts in community centroid to be distinguished from differences in within-group spread. Community structure was rendered using non-metric multidimensional scaling (NMDS; k = 2, trymax = 100), and the fifteen most abundant genera were projected onto the ordination via envfit (999 permutations) to flag taxa aligned with the principal axes.

### Constrained ordination

To quantify the proportion of compositional variation explained by predictors, canonical analysis of principal coordinates (CAP) was implemented through capscale (vegan). Independent models were constructed for the Bray-Curtis and Jaccard matrices, each constrained by *Chlamydiota* status, sex, and husbandry facility. Overall model significance was evaluated by 999 permutations, and per-term contributions were obtained sequentially through anova.cca with by = "term"; adjusted R^2^ was reported to penalise predictor count relative to sample size. Samples were displayed on the first two constrained axes (CAP1 and CAP2), with the constraining variables rendered as biplot arrows whose orientation and length convey their association with the constrained space. To highlight taxonomic contributors to the partitioned variance, genus-level abundances were post-fitted as environmental vectors using envfit (999 permutations); the fifteen genera with the largest correlation coefficients (r^2^) were retained and superimposed on the ordination. Because by = "term" testing is sequential and therefore order-dependent, these results were interpreted alongside the marginal PERMANOVA framework described above.

### Taxonomic composition and differential abundance taxa testing

To provide an overview of cloacal microbiota composition in the presence and absence of *Chlamydiota*, taxonomic profiles were summarised at phylum (top 10) and genus (top 25) level prior to differential abundance testing.

Differential abundance testing of bacterial genera in the presence and absence of *Chlamydiota* was performed on non-rarefied count data using Analysis of Compositions of Microbiome with Bias Correction (ANCOM-BC2) in R. Reads aggregated to genus level were modelled with *Chlamydiota* status, sex, and husbandry facility as fixed effects, using *Chlamydiota*-absent (negative) as reference. P-values were adjusted using Benjamini-Hochberg (BH) False Discovery Rate (FDR) correction (q < 0.05), and biologically meaningful differences were defined as an absolute log2FC > 1.

### Multivariate feature selection and identification of Chlamydiota-associated taxa

We employed two multivariate approaches to explore associations between genus-level composition and *Chlamydiota* status. Both methods used genus-level aggregated data as described above. Abundance data were centred log-ratio (CLR) transformed prior to analysis to address compositional constraints. Zeros were replaced prior to CLR transformation using the multiplicative replacement strategy [43,44], substituting each zero with a small value below the smallest observed non-zero count within that sample.

For elastic net regularised logistic regression, host sex and husbandry facility were incorporated as unpenalised covariates in the model matrix, ensuring these confounders were retained regardless of regularisation strength. The elastic net mixing parameter (𝛼 = 0.5) balanced L1 and L2 penalties to handle correlated genera. The optimal regularisation parameter (𝜆) was selected by 10-fold cross-validation minimising binomial deviance, the standard criterion for penalised logistic regression. As area under the receiver operating characteristic curve (AUC) cannot be reliably estimated with fewer than approximately ten observations per fold at this sample size, model discrimination was evaluated separately by 5-fold cross-validation using AUC, with a 95% confidence interval derived from the cross-validation standard error; the apparent (training-set) AUC is additionally reported to quantify optimism. Feature stability was assessed through 999 bootstrap iterations; genera selected in >60% of resampled datasets were considered stable, consistent with the stability selection framework of Meinshausen and Bühlmann [45], that recommend threshold in the range of 0.6–0.9. As a sensitivity analysis, feature selection and stability were additionally evaluated at the more conservative 𝜆_1se. Selected genera were characterised by their regression coefficients, fold changes, prevalence, and directional association with *Chlamydiota*.

For sparse partial least squares discriminant analysis (sPLS-DA), confounder effects were removed via residualisation prior to modelling, wherein for each genus, we fitted a linear model with sex and husbandry as predictors (excluding *Chlamydiota* status), and used the residuals as confounder-adjusted abundances. The optimal number of components and features per component were determined through five-fold cross-validation repeated 100 times, minimising the balanced error rate with centroids distance. Folds were stratified by *Chlamydiota* status, the default behaviour in mixOmics [46], ensuring balanced class representation across folds. Selected genera were visualised via heatmap (ComplexHeatmap), with confounder-residualised CLR values standardised to Z-scores per genus, genera were pooled across all retained discriminant components, samples ordered by *Chlamydiota* status, and genera hierarchically clustered using Pearson correlation distance with Ward’s D2 linkage. Bar annotations denoted mean scaled abundance per group.

### SparCC network analysis

Genus-level co-occurrence networks were constructed for the presence and absence of *Chlamydiota* through the NetCoMi package to delineate microbial association patterns and identify candidate interaction modules within the cloacal community. The genus pool was restricted to taxa meeting the same within-group prevalence and abundance thresholds used in the beta diversity analysis. Prior to network inference, zero counts in the resulting genus matrix were handled using the pseudoZO option of netConstruct(), whilst non-zero counts were left unchanged. Pairwise genus associations were inferred with SparCC using 200 outer and 100 inner iterations, and edges were retained only where the absolute correlation exceeded 0.45 and the local false discovery rate (lfdr) fell below 0.05. Topological characterisation was performed independently for each network at both the global and nodal scales. Global descriptors comprised node and edge counts, density, modularity, clustering coefficient, average path length, and the percentage of positive edges, whilst nodal centrality was captured by degree, betweenness, closeness, and eigenvector measures, each rescaled via a log-normal fit to allow comparability across networks. Module structure was recovered using the fast-greedy clustering algorithm, and a stringent hub criterion was applied whereby a taxon was classified as a hub only if it simultaneously surpassed the 90th percentile in degree, eigenvector, and betweenness centralities. Group-level structural divergence was then assessed through permutation testing (100 permutations), with adaptive BH correction applied to control the false discovery rate across the multiple network statistics tested. To quantify between-group similarity, the Jaccard index was computed for the overlap of hub sets and the Adjusted Rand Index was used to measure concordance of the recovered cluster assignments.

### PICRUSt2 functional analysis

To infer functional capacity from 16S rRNA gene amplicon data, ASV-level count tables were filtered using the same within-group prevalence and abundance thresholds used in the beta diversity analysis. Filtered ASV sequences and count tables were used as input to PICRUSt2 v2.5.2, which infers metagenome composition based on phylogenetic placement of marker gene sequences against a reference genome database. Predicted KEGG Ortholog (KO) abundances, Enzyme Commission (EC) number abundances, MetaCyc pathway abundances, and taxon-stratified KO contributions were generated with stratified output and per-sequence contribution tracking enabled. Samples with a weighted nearest sequenced taxon index (NSTI) > 0.2 were excluded from downstream analysis to ensure prediction reliability. Pathway abundances were further filtered to retain only those present in at least 60% of samples in at least one sample group. Differential pathway abundance was assessed using two approaches. First, ALDEx2 was applied as a compositionally aware method employing Dirichlet Monte Carlo sampling, Welch’s t-test and BH correction. Second, bootstrap-stabilised elastic net regularised logistic regression (𝛼 = 0.5) was applied over 100 bootstrap iterations, retaining only features selected in ≥ 60% of bootstrap models as a stability criterion [45]. Husbandry facility and host sex were included as unpenalised covariates. For ALDEx2, significance was defined by combined criteria of adjusted p < 0.05, effect size ≥ 1.0, and CLR difference > 0.5; for elastic net, pathways were retained if they showed an absolute CLR difference > 0.5 and were selected in ≥ 60% of bootstrap iterations.

For each significant pathway, constituent KOs were identified through a hierarchical mapping approach. MetaCyc pathway definitions were first decomposed into their constituent reactions and associated EC numbers, which were then mapped to KOs using the curated KEGG REST API. EC-level differential abundance was assessed using ALDEx2 (prevalence ≥ 10%, applying the same significance criteria as described above with an absolute CLR difference > 0.5). Only differentially abundant ECs were retained for downstream KO mapping, ensuring functional consistency between pathway- and gene-level signals. KO specificity was further refined by retaining only those exhibiting a Spearman correlation ≥ 0.3 with the corresponding pathway abundance across samples. KO-level differential abundance was then assessed using ALDEx2 with relaxed thresholds (prevalence ≥ 10%, CLR difference > 0.5, effect size ≥ 1.0). Taxon-level contributions to each KO were derived from PICRUSt2 stratified table, normalised within each sample by total KO abundance and aggregated at the genus level. Results were visualised using multi-level network and bubble plots linking pathways, ECs, KOs, and contributing taxa. Pathway importance was summarised using a composite score integrating five components: (1) perturbation rate, defined as the proportion of a pathway’s detected KOs reaching differential abundance thresholds; (2) the proportion of significant KOs relative to those detected; (3) directional coherence, measuring the consistency of up- or down-regulation across significant KOs; (4) mean effect size across pathway KOs; and (5) KO coverage, defined as the proportion of database-defined pathway genes detected in the dataset. Components were combined into a scaled composite score to rank pathways by overall functional signal strength.

### Characterisation of Chlamydiota-assigned ASVs and BLASTn verification

To characterise *Chlamydiota*-assigned sequences within the dataset, ASVs classified at the phylum level as *Chlamydiota* was extracted from the phyloseq object using the Greengenes2-derived taxonomy table. For each identified ASV, the corresponding representative sequence was retrieved from the DADA2-generated FASTA file of denoised sequences. Taxonomic metadata (genus, species, and classifier confidence) were compiled into a structured output table. To verify and refine taxonomic assignments, each unique *Chlamydiota* ASV sequence was submitted to NCBI BLASTn against the NCBI 16S ribosomal RNA database. The top hit identity percentage and matched species were recorded for each ASV.

## Results

### Study samples and quality assessment

Of the 70 samples retained after rarefaction, 18 (25.7%) belonged to the *Henophidia* clade and 52 (74.3%) to the *Caenophidia* clade. Because we have previously shown that cloacal microbiota composition differs substantially between these two clades [47], we initially considered a stratified analysis encompassing both groups. However, the highly unbalanced distribution of *Chlamydiota*-positive (n = 17) and *Chlamydiota*-negative (n = 1) samples within the *Henophidia* clade precluded meaningful statistical comparison. Hence, all downstream microbiota analyses were restricted to the 52 *Caenophidia* samples. For completeness, taxonomic assignment of *Chlamydiota*-associated ASVs at the genus and species level is nonetheless reported for all 99 samples. The 52 caenophidian snakes originated from four private facilities, and comprised 25 females and 27 males (Fig. 1; Table S2).

**Fig. 1.**
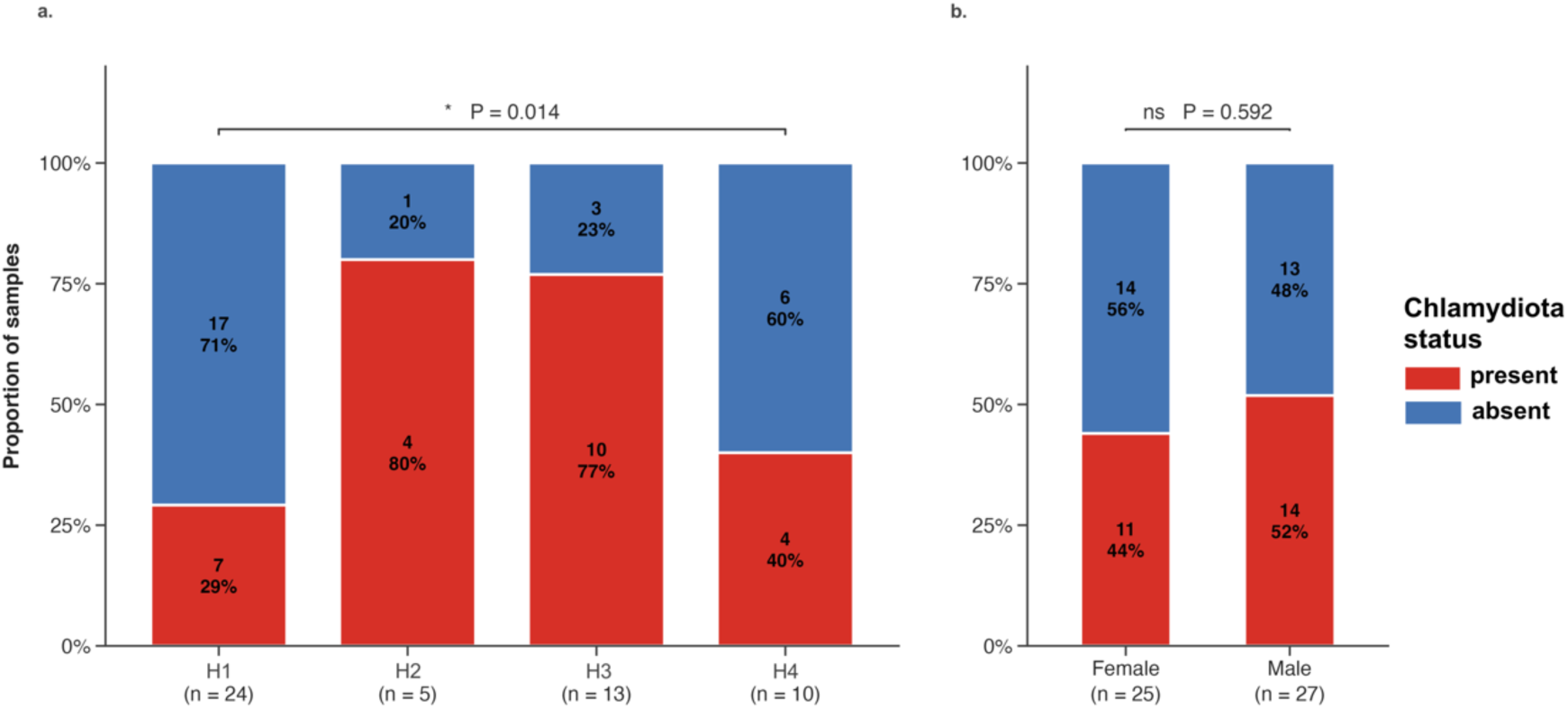
Distribution of *Chlamydiota* in snake cloaca across husbandry facility and sex. Stacked bars show the proportion of *Chlamydiota*-positive and *Chlamydiota*-negative snakes within each category. Absolute sample counts and within-category percentages are displayed inside each segment, and the total number of samples per category is reported below the corresponding axis label (n). (a) *Chlamydiota* status across the four husbandry facilities included in the study. (b) *Chlamydiota* status between female and male snakes. Statistical significance was assessed by Fisher’s exact test under the null hypothesis that *Chlamydiota* status is independent of husbandry facility (a) or sex (b); a significant result indicates a non-random association between *Chlamydiota* status and the categorical variable.

### Bacterial diversity does not differ in the presence and absence of Chlamydiota

A total of 997 ASVs were detected across all samples, with a mean of 104.5 observed ASVs per sample (Table S1). Alpha diversity was assessed using three metrics: Chao1, Shannon, and Pielou’s evenness. After accounting for confounding variables (host sex and husbandry facility), no significant difference in bacterial richness was observed in the presence and absence of *Chlamydiota* (Chao1: F = 0.275, p = 0.602), with estimated marginal means of 66.3 ± 5.7 and 61.7 ± 6.8, respectively (Fig. 2a). Similarly, no significant differences were observed between groups for Shannon diversity (infected: 2.19 ± 0.13; uninfected: 2.17 ± 0.16; F = 0.009, p = 0.924) or Pielou’s evenness (infected: 0.545 ± 0.028; uninfected: 0.554 ± 0.033; F = 0.044, p = 0.835) (Fig. 2a).

**Fig. 2.**
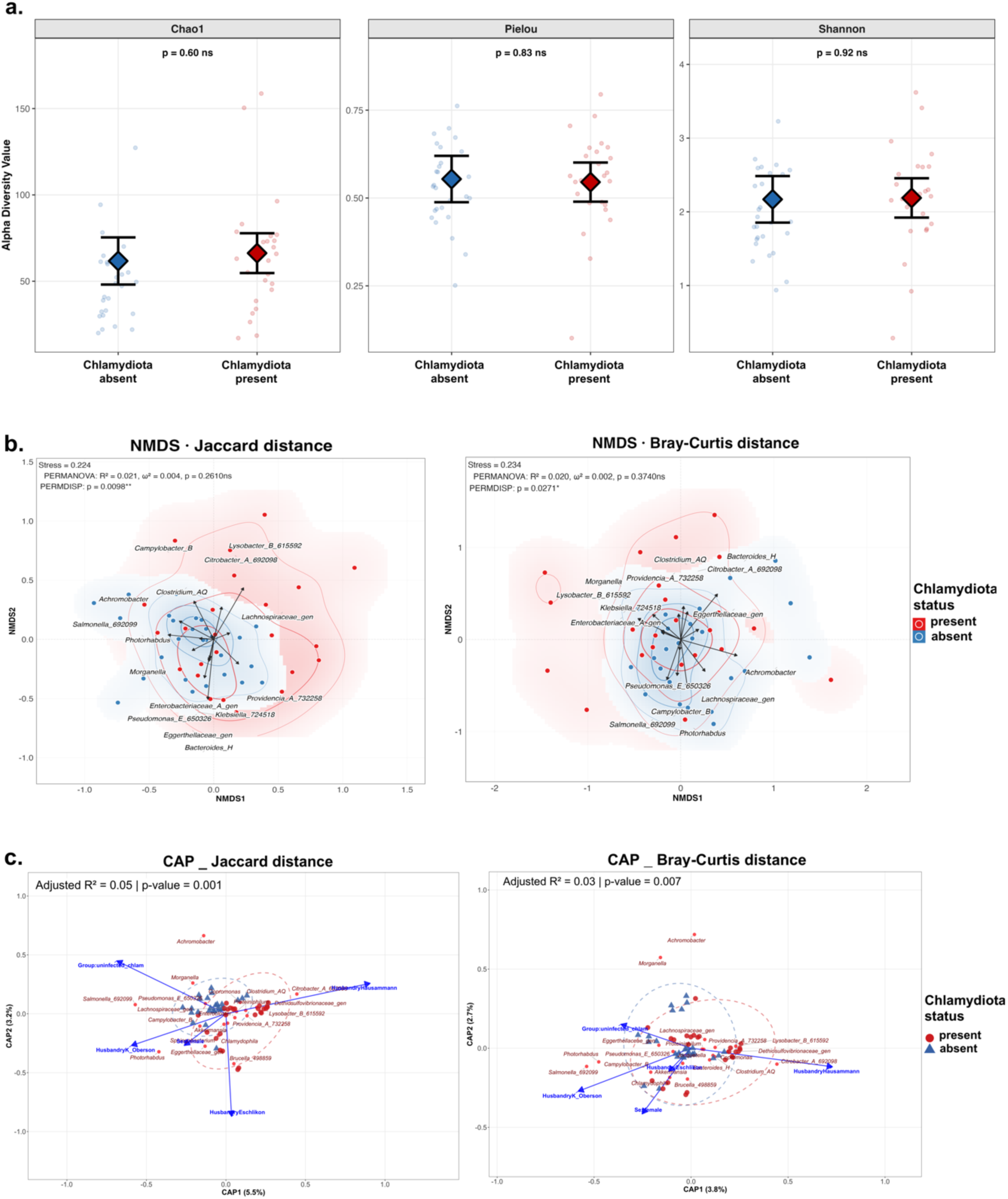
Alpha and beta diversity of cloacal microbiota in relation to the presence and absence of *Chlamydiota*. (a) Estimated marginal means (±95% confidence intervals) for three alpha diversity metrics (calculated from ASV-level data) including Chao1 bacterial richness, Pielou’s evenness, and Shannon diversity, adjusted for host sex and husbandry facility. Large diamonds represent estimated marginal means, error bars indicate 95% confidence intervals, and individual sample values are shown as semi-transparent points. (b) NMDS ordinations at genus level using Jaccard and Bray-Curtis distance metrics. Each point represents an individual snake sample, coloured by *Chlamydiota* status. Shaded density regions and overlaid contour lines illustrate the distributional spread of each group within ordination space. Stress values quantify the goodness-of-fit between the original multidimensional distances and their two-dimensional representation. PERMANOVA R^2^ values represent the proportion of compositional variance explained by *Chlamydiota* status after accounting for sex and husbandry facility effects; omega-squared (𝜔^2^) values indicate effect sizes accounting for unequal sample sizes. PERMDISP p-values indicate whether compositional dispersion differs significantly between *Chlamydiota*-positive and negative groups. Genera vectors (black arrows) indicate the direction and magnitude of association between the 15 most strongly correlated genera and the ordination space; arrow direction shows the gradient of increasing relative abundance for that genus, whilst arrow length reflects the strength of correlation with the ordination axes. (c) CAP ordination of cloacal microbiota composition based on Bray-Curtis and Jaccard distance metrics. Each point represents an individual snake sample, coloured by *Chlamydiota* status. Arrows represent fitted environmental variables, with direction indicating the gradients of variation and length reflecting the strength of association with the ordination axes. Red points indicate the top 15 genera most strongly correlated with the ordination axes. Dashed ellipses enclose samples from each *Chlamydiota* status group.

### Chlamydiota status is not associated with major shifts in cloacal microbiota composition

PERMANOVA analysis of genus-level community composition revealed no significant differences between *Chlamydiota*-positive and negative snakes after adjustment for host sex and husbandry facility, with *Chlamydiota* status explaining only a small proportion of variance in both Bray-Curtis (p = 0.374, 𝜔^2^ = 0.002) and Jaccard (p = 0.261, 𝜔^2^ = 0.004) distance matrices (Fig. 2b; Table S3_1). Husbandry facility was the dominant contributor to microbiota variation, accounting for 8.3% and 10.8% of Bray-Curtis and Jaccard variance, respectively, whereas host sex was not significantly associated with community composition. NMDS ordination confirmed overlap between *Chlamydiota*-positive and negative groups (Fig. 2b).

Despite the absence of significant centroid separation, PERMDISP analysis identified significantly higher within-group compositional heterogeneity in the presence of *Chlamydiota* for both Bray-Curtis (mean difference = 0.075, 95% CI: 0.009–0.141, p = 0.027) and Jaccard (mean difference = 0.065, 95% CI: 0.016–0.113, p = 0.010) distances (Fig. 2b; Table S3_2).

Environmental vector fitting identified several genera correlated with ordination structure. For Bray-Curtis distance, the strongest associations were with *Bacteroides*_H (R^2^ = 0.40), *Achromobacter* (R^2^ = 0.40), *Photorhabdus* (R^2^ = 0.27), *Lysobacter*_B (R^2^ = 0.26), and *Salmonella* (R^2^ = 0.21), with additional contributions from *Lachnospiraceae*_gen (unclassified members of *Lachnospiraceae* family), *Morganella*, *Campylobacter*_B, and *Clostridium*_AQ. For Jaccard distance, *Bacteroides*_H and *Lysobacter*_B were the strongest contributors, with additional associations from *Salmonella* and *Eggerthellaceae*_gen (Fig. 2b; Table S3_3).

### Chlamydiota presence contributes to cloacal microbial community variation

To quantify how much of the variation in cloacal community composition was explained by *Chlamydiota* status, host sex, and husbandry facility, we used CAP with these variables as constraints. For Jaccard distance, the full model explained 14.5% of variance (adjusted r^2^ = 0.05, p = 0.001) (Fig. 2c), with husbandry facility contributing 8.60% (p = 0.001), *Chlamydiota* status 3.50% (p = 0.004), and sex 2.39% (p = 0.061) (Table S4_1). For Bray-Curtis distance, the full constrained model explained 12.3% of total variance (adjusted r^2^ = 0.03, p = 0.007) (Fig. 2c), with husbandry facility contributing 7.59% (p = 0.015), sex 2.35% (p = 0.138), and *Chlamydiota* status 2.33% (p = 0.174) (Table S4_2). Given the reported confounding between *Chlamydiota* status and husbandry facility (Fisher’s exact test, p = 0.015), the significant Jaccard effect for *Chlamydiota* should be interpreted cautiously. A sensitivity analysis incorporating an interaction term between *Chlamydiota* status and husbandry facility found no evidence that the effect of *Chlamydiota* on community composition varied across facilities (Jaccard: F = 0.92, p = 0.752; Bray–Curtis: F = 0.89, p = 0.761), supporting the use of the main-effects model presented here.

Post-hoc envfit identified genera whose abundance correlated with the constrained ordination axes, most strongly *Achromobacter* under both Jaccard (vector length = 0.68) and Bray-Curtis (0.72) distances, followed at lower vector lengths by *Salmonella*, *Photorhabdus*, *Citrobacter*_A, *Morganella*, and *Lysobacter*_B (shared between both distances), with *Brucella* and *Eggerthellaceae*_gen additionally identified under Jaccard, and *Campylobacter*_B and *Chlamydophila* under Bray-Curtis (Fig. 2c; Table S4_3).

### Specific taxa may distinguish presence and absence of Chlamydiota in snake cloaca

At the phylum level, both groups were dominated on average by *Proteobacteria*, followed by *Bacteroidota*, *Actinobacteriota*, and *Firmicutes*_A, with *Synergistota* enriched in the presence of *Chlamydiota*. At the genus level, *Salmonella*, *Bacteroides*_H, and *Morganella* were the most abundant genera across both groups, with several less abundant taxa showing group-specific enrichment (Fig. S1).

Because mean relative abundances and presence/absence patterns are descriptive and may be confounded, we used ANCOM-BC2 to test for differential abundance between the groups on the unrarefied count data, adjusting for sex and husbandry facility. After multiple testing correction and application of an effect size threshold (log2FC > 1), ten genera differed significantly between groups (q < 0.05), all of which exceeded the effect size threshold. Five were enriched in the presence of *Chlamydiota*, most prominently an *Neisseriaceae*_gen (log2FC = +2.40, q = 0.014) and *Eggerthella* (log2FC = +2.28, q = 0.014), followed by *Proteus* (log2FC = +1.64, q = 0.014), *Desulfovibrio* (log2FC = +1.25, q = 0.014), and an *Chitinophagaceae*_gen (log2FC = +1.35, q = 0.040), and five were enriched in the absence of *Chlamydiota*: *Stenotrophomonas* (log2FC = −1.99, q = 0.029), *Photorhabdus* (log2FC = −1.96, q = 0.020), *Lachnospiraceae*_gen (log2FC = −1.82, q = 0.040), *Copromonas* (log2FC = −1.58, q = 0.040), and *Brucella* (log2FC = −1.45, q = 0.029) (Fig. 3a). Three additional genera (*Brevibacterium*, *Brevilactibacter*, and *Paracoccus*) were nominally significant (p < 0.05) but did not survive multiple testing correction and did not meet the effect size threshold (Fig. 3b).

**Fig. 3.**
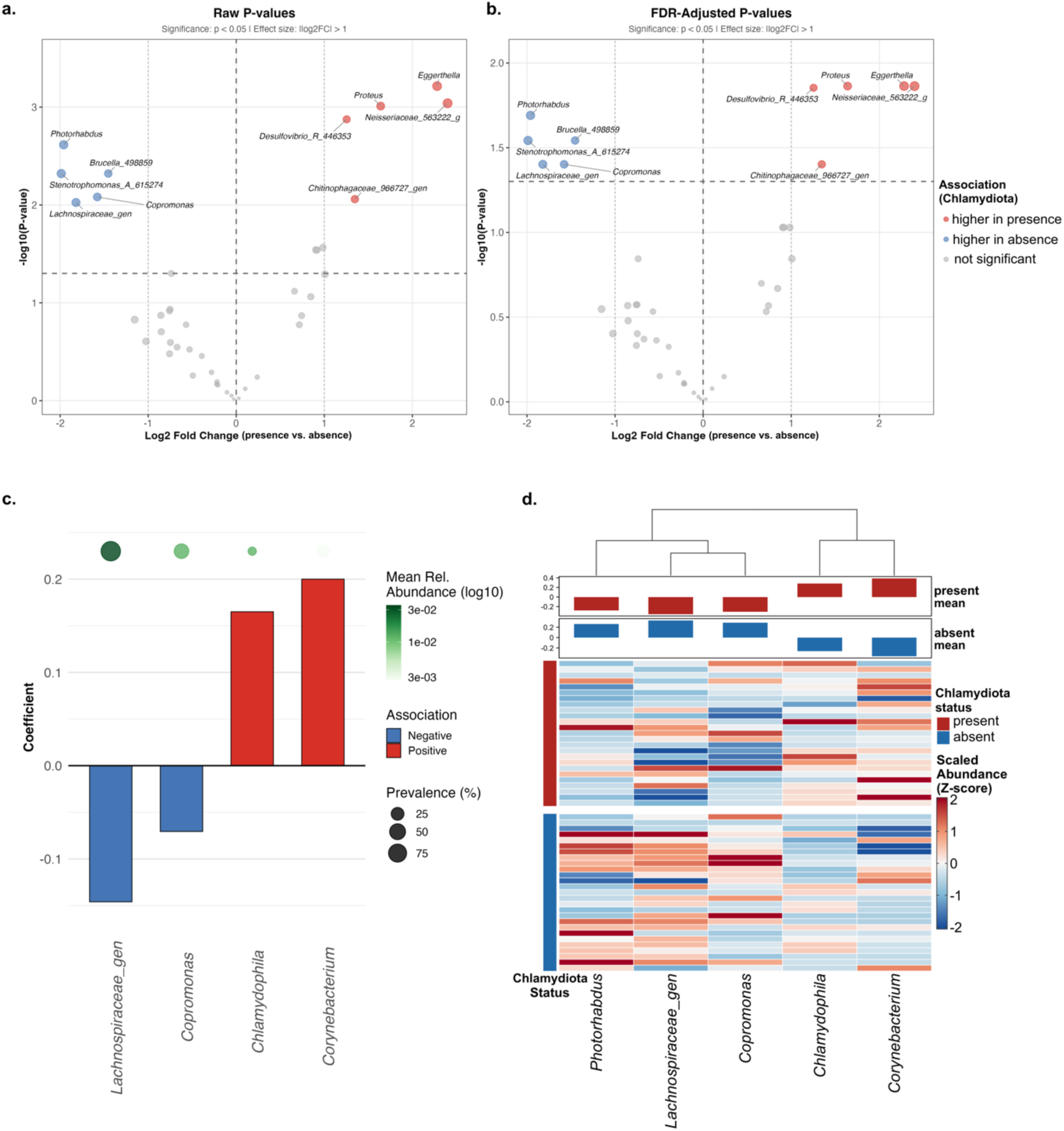
Differential abundance and multivariate identification of microbial genera in the presence and absence of *Chlamydiota* in snake cloaca. (a–b) Volcano plots displaying the relationship between effect size (log2FC, *Chlamydiota*-positive *vs*. negative) and statistical significance for genus-level differential abundance testing using ANCOM-BC2. Panel (a) shows raw p-values and panel (b) shows BH-corrected p-values, both expressed as -log10 transformed values. Each point represents a genus, coloured by direction and significance (red: significantly higher in the presence, log2FC > 1, p < 0.05; blue: significantly higher in absence of *Chlamydiota*, log2FC > 1, p < 0.05; grey: not significant or below effect size threshold), with point size proportional to log2FC. The horizontal dashed line denotes the significance threshold (p = 0.05) and vertical dotted lines indicate the log2FC = 1 effect size threshold. (c) Genera selected by elastic net regularised logistic regression (α = 0.5) after controlling for sex and husbandry facility, restricted to genera meeting the stability criterion (selected in > 60% of 999 bootstrap resamples at 𝜆_min). Bar height represents the elastic net coefficient magnitude, with bar colour indicating direction of association (red = enriched in the presence; blue = enriched in the absence of *Chlamydiota*). Bubbles positioned above bars indicate overall sample prevalence (size) and mean relative abundance (colour, log10 scale) for each genus. (d) Heatmap of CLR-transformed, confounder-residualised abundances for genera selected across all sPLS-DA components, standardised to Z-scores column-wise. Samples are grouped by *Chlamydiota* status and genera are hierarchically clustered using Pearson correlation distance with Ward’s linkage, with the dendrogram shown above. Bar annotations indicate mean scaled abundance within *Chlamydiota*-positive (red) and negative (blue) groups.

### Multivariate models reveal genera discriminating Chlamydiota-positive from negative microbiota

Elastic net regularised logistic regression, a predictive classifier that simultaneously selects and weights genera to discriminate *Chlamydiota*-positive from negative samples, identified 12 genera as features in the fitted model (Table S5_1). Of these, four met stability criteria (> 60% bootstrap selection frequency across 999 iterations): *Corynebacterium*, positively associated with the presence (coefficient = 0.200, 73.4% selection frequency, 2.3-fold enrichment), and two genera negatively associated with the presence of *Chlamydiota*, *Lachnospiraceae*_gen (coefficient = -0.146, 79.4% selection frequency) and *Copromonas* (coefficient = -0.071, 60.7% selection frequency) (Fig. 3c; Table S5_2). Model discrimination was moderate when evaluated by cross-validation (CV AUC = 0.74, 95% CI: 0.58-0.90; apparent training AUC = 0.90, reflecting the optimism expected for penalised models at small sample sizes). A sensitivity analysis at the more conservative 𝜆_1se retained a single genus and yielded essentially unchanged cross-validated discrimination (AUC = 0.74), indicating that the moderate performance is not an artefact of the 𝜆_min choice.

Through an alternative analytical framework, sPLS-DA after confounder residualisation identified a single discriminant component selecting five genera (model accuracy 46.5%) (Fig. 3d; Table S6_1). The lower balanced accuracy of this model (46.5%) compared with the elastic net (cross-validated AUC = 0.74) can reflect the two methods’ differing treatment of confounders: the elastic net retained sex and husbandry as unpenalised covariates within the model, whereas the sPLS-DA was applied to abundances from which these effects had first been residualised, a more conservative step that removes any *Chlamydiota*-associated variance correlated with husbandry or sex. *Corynebacterium* exhibited the strongest positive association with *Chlamydiota* (loading = 0.695), whilst *Lachnospiraceae*_gen showed the strongest negative association (loading = -0.556), followed by *Copromonas* (loading = -0.336), and *Photorhabdus* (loading = -0.216) (Table S7_2). Hierarchical clustering of these genera based on CLR-transformed, confounder-residualised abundance profiles revealed two distinct co-abundance patterns: *Corynebacterium* consistently displayed elevated scaled abundances in the presence of *Chlamydiota*, whilst *Lachnospiraceae*_gen, *Copromonas*, and *Photorhabdus* exhibited relatively higher abundances in the absence of *Chlamydiota* (Fig. 3d; Table S6_2).

### A cross-phylum anaerobic guild anchors the co-occurrence network associated with the presence of *Chlamydiota*

To assess whether microbial association patterns varied with *Chlamydiota* status, SparCC was applied to each group to infer genus-level co-occurrence networks. Both networks were sparse (uninfected: 14 edges, density = 0.005; infected: 34 edges, density = 0.012), but differed in the number and type of the associations recovered (Fig. 4; Table S7_1). The network in the absence of *Chlamydiota* contained fewer edges, a notable proportion of which were negative (28.6%; SparCC r = -0.622 to 0.769), whereas in the presence of *Chlamydiota* the network harboured 2.4-fold more edges, the large majority of which were positive (97.1%; SparCC r = -0.529 to 0.712). Topological metrics also differed, modularity (0.86 *vs*. 0.59) and clustering coefficient (0.64 *vs*. 0.50) were higher in the presence of *Chlamydiota*.

**Fig. 4.**
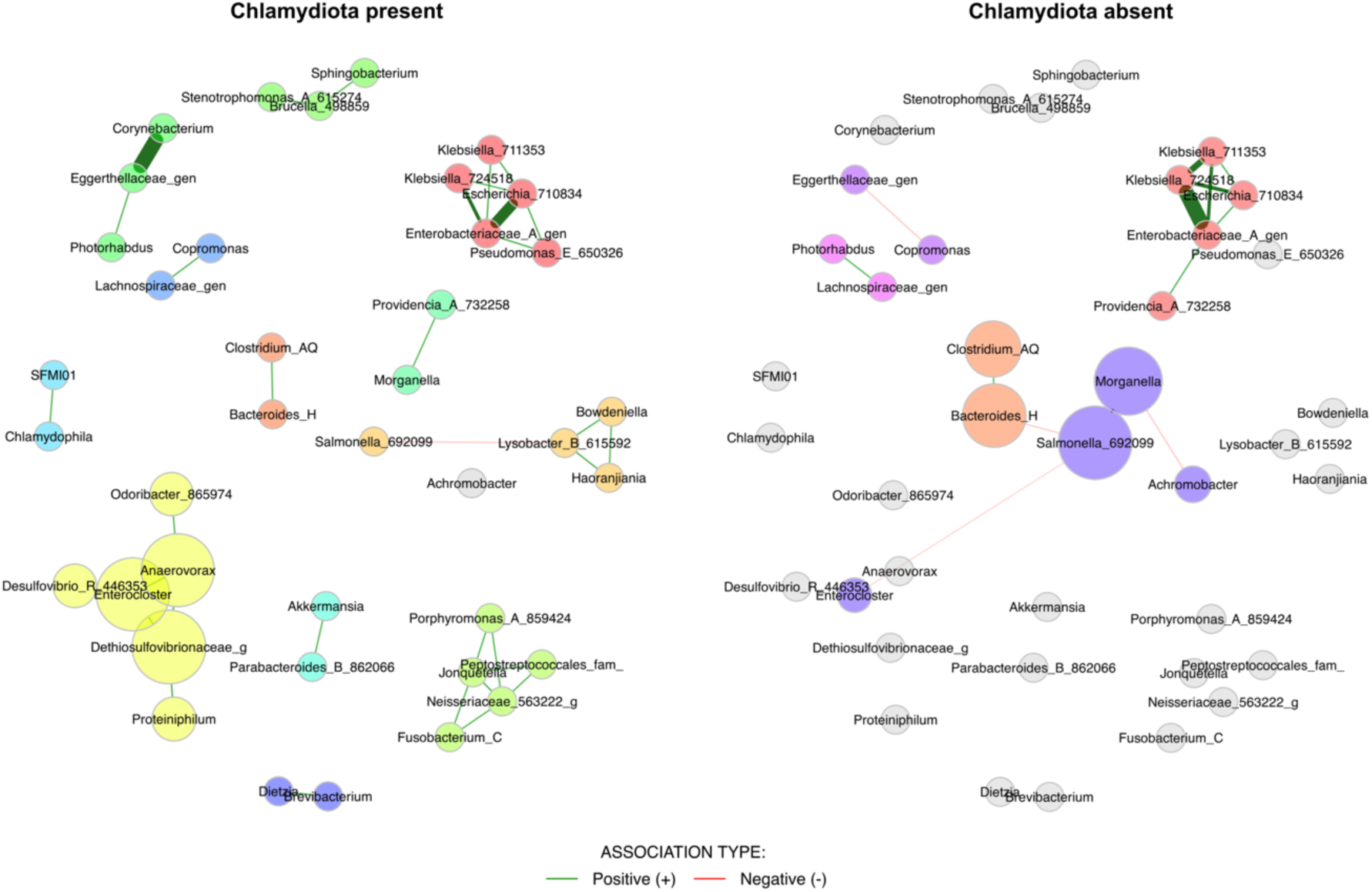
Genus-level co-occurrence networks for cloacal microbiota in the presence and absence of *Chlamydiota*. Association inference was performed using SparCC, with networks built independently for each infection group. Each node corresponds to a genus, with node size proportional to weighted degree centrality. Community structure within each network was identified through fast-greedy modularity optimisation, so node colours denote within-network module membership and are not comparable between panels. Edges depict statistically supported associations (r > 0.45, lfdr < 0.05); edge opacity scales with association strength, and edge colour indicates association direction (green: positive; red: negative). Genera lacking edges in both networks were omitted from the visualisation.

In the absence of *Chlamydiota*, the recovered positive associations were dominated by *Enterobacteriaceae*, with eight of the ten positive edges involving members of this family (e.g. *Klebsiella*, *Enterobacteriaceae*_gen, *Escherichia*, *Salmonella*); the most central nodes were *Salmonella*, *Morganella*, and *Bacteroides*_H (Fig. 4; Tables S7_2 and S7_3). The four negative associations were *Enterocloster*-*Salmonella* (r = -0.622), *Bacteroides*_H-*Salmonella* (r = -0.502), *Achromobacter*-*Morganella* (r = -0.494), and *Copromonas*-*Eggerthellaceae*_gen (r = -0.472). In the presence of *Chlamydiota*, the largest connected module formed a cross-phylum anaerobic guild anchored by unclassified *Dethiosulfovibrionaceae*, *Enterocloster*, and *Anaerovorax*, extending to *Proteiniphilum* and *Desulfovibrio* and bridging *Synergistota*, *Firmicutes*, *Bacteroidota*, and *Desulfobacterota* (Fig. 4; Tables S7_2 and S7_3). Other recovered associations included a *Corynebacterium*-*Eggerthellaceae*_gen edge (r = 0.712; both *Actinobacteriota*) and a *Gammaproteobacteria* sub-cluster encompassing *Enterobacteriaceae*_gen, *Escherichia*, *Klebsiella*, and *Pseudomonas*_E (r = 0.567-0.677). Because *Chlamydophila* was detected only in positive samples by definition, its associations could be assessed solely within the network in the presence of *Chlamydiota*, where it formed a single weak positive edge with *Christensenellaceae*_gen (r = 0.489) (Fig. 4; Table S7_1).

### Inferred functional differences between the cloacal microbiota in the presence and absence of Chlamydiota are subtle

Functional potential was predicted from 16S rRNA gene data using PICRUSt2 and analysed at both KEGG (376 pathways) and MetaCyc (452 pathways) levels. PERMANOVA on Bray-Curtis distances revealed no significant effect of *Chlamydiota* status on functional composition at either level (KEGG: R^2^ = 0.011, p = 0.566; MetaCyc: R^2^ = 0.018, p = 0.396), with homogeneous within-group dispersion (Table S8_1 and S8_2), and ALDEx2 compositional testing identified no significantly differentially abundant pathway in either database. These analyses indicate no statistically significant functional divergence between groups.

Bootstrap-stabilised elastic net regression was used to identify pathways with predictive value, recovering no KEGG and three MetaCyc pathways at the ≥ 60% selection-frequency threshold (Fig. 5; Table S8_3-5). All three, mycolyl-arabinogalactan-peptidoglycan complex, mono-trans, poly-cis decaprenyl phosphate, and mycothiol biosynthesis, were predictive of *Chlamydiota* presence and converged on a coherent actinobacterial signature, being taxonomically anchored by *Dietzia*, *Corynebacterium*, and *Brevibacterium*/*Brevilactibacter*, the same actinobacterial genera independently enriched in the presence of *Chlamydiota* (Fig. 5; Table S8_4). The inferred functional signal therefore reflects this underlying taxonomic enrichment rather than independent functional reprogramming.

**Fig. 5.**
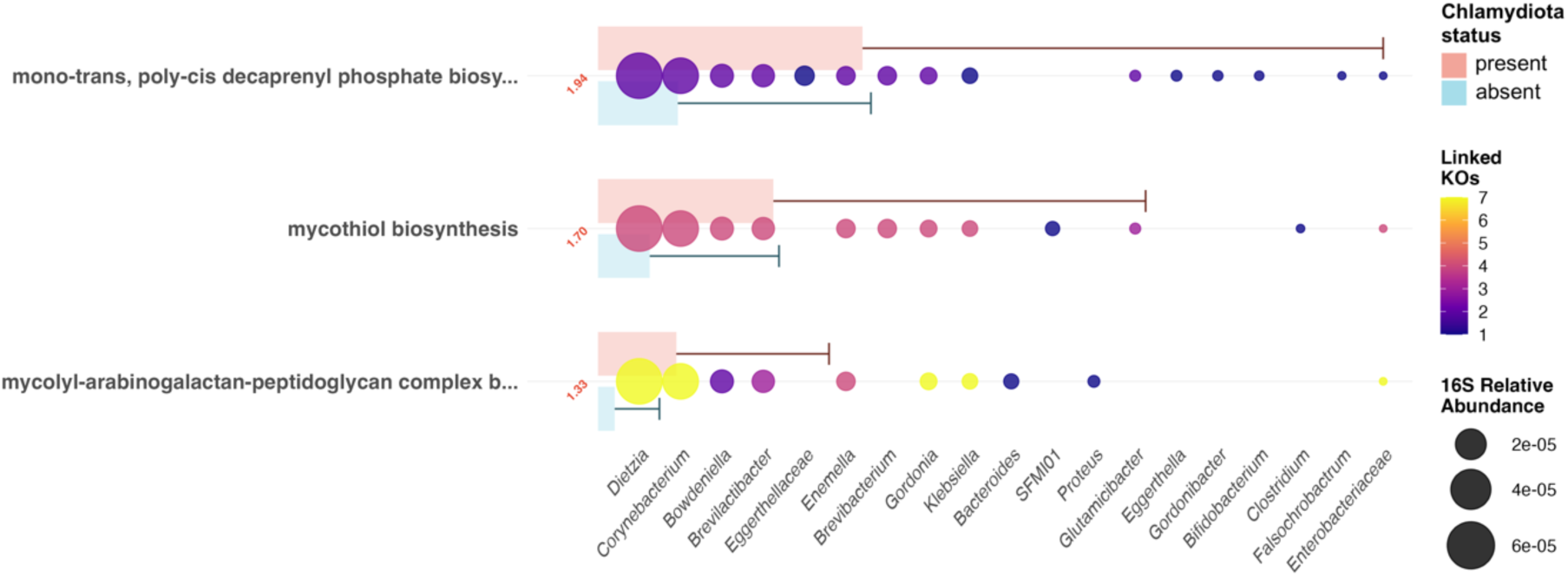
Stable elastic net-selected pathways and their taxon-level functional linkages from the MetaCyc analysis. Pathways are ranked by CLR difference between the presence and absence of *Chlamydiota*, with CLR difference values shown adjacent to the y-axis and colour-coded by enrichment direction. Transparent horizontal bars indicate mean predicted pathway abundance (± SD) per group. Genera (x-axis, italic) represent taxa contributing to the displayed pathways through KEGG Ortholog (KO)-mediated linkage, ordered by total abundance. Each bubble at a pathway-taxon intersection denotes a functional connection mediated by one or more KOs; bubble size is proportional to the mean 16S rRNA gene-derived relative abundance of the corresponding genus across all samples, while bubble colour intensity reflects the number of distinct KOs linking that taxon to the pathway, brighter colours indicating broader enzymatic contribution.

### Targeted analysis reveals diverse Chlamydiota lineages in snake cloacal samples

Out of 59 samples initially identified as positive for *Chlamydiota*, 12 tested positive for *Chlamydiota* using both pan-*Chlamydiota* PCR (mean ct = 29, SD = 4.7) and 16S rRNA amplicon sequencing, 43 were positive only by pan-*Chlamydiota* PCR (mean ct = 33.3, SD = 1.4) and 4 were positive only using 16S rRNA amplicon sequencing. In total, at least one *Chlamydiota* ASV was assigned to 16 samples. Taxonomic assignment at family level revealed that the majority of reads belonged to family *Chlamydiaceae* (70%), and a smaller proportion assigned to families *Parachlamydiaceae* (25%) and *Rhabdochlamydiaceae* (5%) (Table 1).

**Table 1.** Sample-level distribution of *Chlamydiota* ASVs with Greengenes2 and NCBI BLASTn taxonomic assignments.

| Sample ID | Husbandry | ASV ID | Genus (greengenes2) | Species (greengenes2) | Confidence† | Rel. abundance (%)‡ | Species (BLASTn) | BLASTn Identities (%)‡ |
| --- | --- | --- | --- | --- | --- | --- | --- | --- |
| S_19444* | H1 | ASV_38 | *Chlamydophila* | *Chlamydophila pneumoniae* | 1 | 0.289 | *Chlamydia serpentis* | 100 |
| S_19453* | H4 | ASV_73 | *Chlamydophila* | *Chlamydophila pneumoniae* | 1 | 0.042 | *Chlamydia serpentis* | 99 |
| S_19455§ | H1 | ASV_2896 | *Neochlamydia* | *Neochlamydia hartmannellae* | 0.6 | 0.001 | *Neochlamydia hartmannellae* | 87 |
| S_19455§ | H1 | ASV_73 | *Chlamydophila* | *Chlamydophila pneumoniae* | 1 | 0.002 | *Chlamydia serpentis* | 99 |
| S_19462# | H3 | ASV_1169 | *Chlamydophila* | *Chlamydophila pneumoniae* | 1 | 0.130 | *Candidatus Chlamydia sanziniae* | 100 |
| S_19547* | H5 | ASV_73 | *Chlamydophila* | *Chlamydophila pneumoniae* | 1 | 49.348 | *Chlamydia serpentis* | 99 |
| S_19556* | H5 | ASV_73 | *Chlamydophila* | *Chlamydophila pneumoniae* | 1 | 0.018 | *Chlamydia serpentis* | 99 |
| S_19559# | H3 | ASV_747 | *Chlamydia* | *Chlamydia muridarum* | 0.68 | 0.886 | *Chlamydia muridarum* | 100 |
| S_19572# | H4 | ASV_1553 | *Neochlamydia* | *Neochlamydia hartmannellae* | 0.83 | 0.077 | *Neochlamydia hartmannellae* | 92 |
| S_19574* | H4 | ASV_38 | *Chlamydophila* | *Chlamydophila pneumoniae* | 1 | 0.093 | *Chlamydia serpentis* | 100 |
| S_19575* | H4 | ASV_38 | *Chlamydophila* | *Chlamydophila pneumoniae* | 1 | 0.063 | *Chlamydia serpentis* | 100 |
| S_19576§ | H4 | ASV_38 | *Chlamydophila* | *Chlamydophila pneumoniae* | 1 | 0.032 | *Chlamydia serpentis* | 100 |
| S_19576# | H4 | ASV_1553 | *Neochlamydia* | *Neochlamydia hartmannellae* | 0.83 | 0.032 | *Neochlamydia hartmannellae* | 92 |
| S_19576# | H4 | ASV_2658 | *Protochlamydia* | *Protochlamydia_sp* | 0.83 | 0.008 | *Candidatus Protochlamydia sp.* | 96 |
| S_19577# | H3 | ASV_38 | *Chlamydophila* | *Chlamydophila pneumoniae* | 1 | 0.131 | *Chlamydia serpentis* | 100 |
| S_19579# | H3 | ASV_38 | *Chlamydophila* | *Chlamydophila pneumoniae* | 1 | 3.668 | *Chlamydia serpentis* | 100 |
| S_19585§ | H3 | ASV_3017 | *Rhabdochlamydiaceae_gen* | *Rhabdochlamydiaceae_gen* | 0.69 | 0.000 | *Candidatus Rhabdochlamydiaceae sp.* | 93 |
| S_19585# | H3 | ASV_38 | *Chlamydophila* | *Chlamydophila pneumoniae* | 1 | 0.540 | *Chlamydia serpentis* | 100 |
| S_19587§ | H3 | ASV_38 | *Chlamydophila* | *Chlamydophila pneumoniae* | 1 | 0.002 | *Chlamydia serpentis* | 100 |
| S_19592§ | H1 | ASV_3033 | *Neochlamydia* | *Neochlamydia_sp* | 0.75 | 0.007 | *Candidatus Protochlamydia naegleriophila* | 94 |

\* Sample from the *Caenophidia* snake clade included in the microbiota analysis.

The dominant ASVs within family *Chlamydiaceae* were ASV_38 (n = 8) and ASV_73 (n = 4), both annotated by the Greengenes2 classifier as *C. pneumoniae* (score = 1). However, BLASTn analysis returned *C. serpentis* as the top hit for both ASVs (99–100% identity). A third *Chlamydiaceae* ASV (ASV_1169), detected in sample S_19462, was annotated as *C. pneumoniae* by Greengenes2 but matched *Candidatus* C. sanziniae at 100% identity by BLASTn. One ASV assigned to *C. muridarum* (ASV_747), detected in sample S_19559, was confirmed as *C. muridarum* by BLASTn (100% identity). Within family *Parachlamydiaceae*, three ASVs were identified. ASV_1553 (*Neochlamydia hartmannellae*, 92% BLASTn identity) was detected in two samples. ASV_2658 (*Candidatus* Protochlamydia sp., 96% identity) was detected in a single sample, and ASV_3033, annotated by Greengenes2 as *Neochlamydia* sp., matched *Candidatus* P. naegleriophila at 94% identity and was detected in one sample. A single ASV belonging to family *Rhabdochlamydiaceae* (ASV_3017; *Candidatus* Rhabdochlamydiaceae sp., 93% identity) was detected in one sample (Table 1).

To characterise *Chlamydiota* carriage levels at the 16S rRNA level, we computed the per-sample relative abundance of all *Chlamydiota*-assigned ASVs on the non-rarefied count data. Across the 16 samples, *Chlamydiota* relative abundance varied widely, from 0.002% to 49.3% of the library (median 0.085%, IQR 0.036-0.352%; Table 1).

## Discussion

Across several studies, chlamydial DNA has been recovered from clinically healthy snakes, with cloacal and choanal shedding [7–9,48]. Conversely, *C. serpentis* has been linked to granulomatous multi-organ disease in captive viperid collections, with intermittent shedding supporting recurrent outbreaks even in previously treated facilities [5,11]. Independent case reports have implicated *C. pneumoniae* in granulomatous splenitis and neurological disease in a royal python and in fatal disseminated disease in emerald tree boas [49,50]. Whilst most *Chlamydiota*-positive samples carried *Chlamydiota* at low relative abundance (median 0.085%), a small subset showed higher carriage, with two *C. serpentis*-assigned samples reaching 3.7% and 49.3% of the library. This wide spectrum of relative carriage levels may correspond to a range of host-microbe states, from low-level detection, which reflects transient passage, environmental contamination, or sub-clinical residency, through to relative abundances compatible with active infection; however, this cannot be established from amplicon data at a single time point. One speculative factor that could contribute to the wide range of *Chlamydiota* relative abundance observed here is hibernation phase. Reptile microbiomes shift substantially across active, pre-hibernation, and hibernation periods, with documented changes in dominant bacterial phyla and metabolism-associated taxa in snakes [51] and broadly similar patterns reported in crocodilians [52]. Hibernation phase was not recorded in the present cohort; whether chlamydial relative abundance varies with seasonal physiology remains an open question that requires longitudinal sampling to test. Placed alongside the mammalian literature, in which *Chlamydia* species behave as long-term gastrointestinal commensals in cattle, sheep, pigs, birds and mice [12,13] yet simultaneously compromise epithelial barrier integrity, goblet cell differentiation and antimicrobial peptide expression in the murine colon [14], and can establish infection in human gastrointestinal organoids accompanied by aberrant developmental forms [53], it appears most parsimonious to suggest snake cloacal *Chlamydiota* as forming a heterogeneous assemblage spanning commensal-like persistence, environmentally acquired *Chlamydia*-like organisms (CLOs), and pathogenic lineages whose clinical manifestation may depend on host species, chlamydial strain, inoculum, and co-infection context.

Within the *Caenophidia* cohort examined here, none of the alpha diversity metrics differed significantly in the presence and absence of *Chlamydiota*, and there were no significant shifts in overall community composition for either Bray-Curtis or Jaccard distances. Whilst we are well aware that the snake cloaca differs substantially from the mammalian intestine and the human cervicovaginal tract in anatomy, physiology, resident microbial baseline, and in the chlamydial species involved [6,7,9,54], the limited body of work available in reptiles leads us to draw broader comparisons with the more established findings in mammals. Bearing this caveat in mind, the modest alpha diversity effects observed here appear to differ from the increase in gut bacterial Shannon diversity reported in *Chlamydia*-infected horses [32] and from the *Lactobacillus*-driven reduction in diversity frequently associated with *C. trachomatis*-positive cervicovaginal microbiomes [27,28,55–57], and seem more consistent with studies in which chlamydial colonisation has been linked to subtle, spatially heterogeneous mucosal remodelling rather than to a uniform microbial community shift [58]. An informative result in our beta diversity analysis was the significant dispersion: *Chlamydiota*-positive samples exhibited markedly greater within-group dispersion than negative samples. An increase in variance without a clear centroid shift may suggest an ecological disturbance and is in line with the Anna Karenina principle in microbial ecology, whereby dysbiotic states are more variable than healthy ones [59–61]. A comparable pattern has been described for *C. trachomatis*-associated cervicovaginal microbiomes, in which community state type IV displays heightened inter-subject variability alongside loss of *Lactobacillus* dominance [28,57]. Our findings suggest that husbandry facility is the dominant predictor in both unconstrained and constrained analyses, in agreement with prior work demonstrating that cage-level environment, cohabitation, diet and hygiene exert a major influence on reptilian gut and cloacal microbiota [20,21]. The confounding between *Chlamydiota* status and facility therefore warrants caution when attributing the modest Jaccard-based CAP effect to *Chlamydiota* status alone.

Taxonomically, the dominance of *Proteobacteria*, followed by *Bacteroidota*, *Actinobacteriota* and *Firmicutes*_A, together with *Salmonella*, *Bacteroides* and *Morganella* as the most abundant genera, is consistent with prior surveys of snake cloacal and faecal microbiota [19–21,62]. ANCOM-BC2, elastic net and sPLS-DA consistently identified *Lachnospiraceae*_gen and *Copromonas* as negatively associated with the presence of *Chlamydiota*. The depletion of *Lachnospiraceae* observed in the presence of *Chlamydiota* is noteworthy, because this family contains major butyrate producers which stabilise tight-junction proteins and reinforces mucosal barrier integrity [63,64]. Reductions in *Lachnospiraceae* are a recurrent feature of mucosal inflammatory and infectious states [63,65], and the parallel depletion of short-chain fatty-acid-producers reported in *Chlamydia*-infected equine gut microbiota [32] suggests that loss of this guild may be a broader signature of chlamydial mucosal colonisation, whilst acknowledging that the ecological baseline of the snake cloaca differs substantially from the mammalian gut. ANCOM-BC2 additionally flagged *Desulfovibrio*, *Brevibacterium*, *Proteus* and *Dietzia* as enriched in the presence of *Chlamydiota*. The enrichment of the *Actinobacteriota* genera *Corynebacterium*, *Dietzia* and *Brevibacterium*, which have been previously recovered from both healthy and diseased snake respiratory surfaces [48], points to an expansion of lipid-rich, cell-wall-specialised taxa whose ecological significance is explored further below in our functional analyses.

The SparCC co-occurrence networks provided an additional, informative perspective on community restructuring. However, with the current sample size, network inference was performed on relatively small groups and therefore the results should be interpreted with caution as exploratory patterns. The network in the presence of *Chlamydiota* was denser, more modular and harboured almost only positive associations, whereas the network in the absence of *Chlamydiota* contained a higher proportion of negative co-occurrence relationships. *Chlamydiota*-positive samples harboured a prominent anaerobic guild, linking bacteria from the *Synergistota*, *Firmicutes*, *Bacteroidota* and *Desulfobacterota* potentially into a tightly cross-feeding module. The expansion of anaerobic taxa alongside chlamydial presence aligns with patterns reported across mammalian mucosal niches, for which both ecological and mechanistic drivers have been proposed. In the cervicovaginal tract, *C. trachomatis* infection is consistently associated with reduced *Lactobacillus* dominance and overgrowth of anaerobic taxa [27,28,55,66,67], and comparable shifts have been reported in the rectal microbiota of men with urogenital *C. trachomatis* infection [56] and in the large intestine of *Chlamydia*-infected horses [32]. Ecologically, chlamydial infection may perturb local oxygen tension and redox balance through host inflammatory responses, conditions that disadvantage aerobic and facultative-anaerobic *Enterobacteriaceae* whilst favouring obligate anaerobes [68,69]. The near-absence of negative edges in the presence of *Chlamydiota* may suggest a shift towards cooperative metabolic dependencies rather than competitive exclusion, a pattern also reported in the gut of *Chlamydia*-infected horses [32] and in the rectal microbiota of *C. trachomatis*-infected men [56,67]. By contrast, the network of *Chlamydiota*-negative samples was dominated by *Enterobacteriaceae*-centred co-occurrences and, unlike the network of positive samples, included several negative associations, between *Salmonella* and *Enterocloster*, *Salmonella* and *Bacteroides*_H, and *Copromonas* and *Eggerthellaceae*_gen. Negative co-occurrence relationships of this kind are previously reported in enteric communities [70,71] and may reflect resource competition, niche partitioning, or host-imposed segregation; however, the underlying mechanism cannot be concluded from cross-sectional co-occurrence data alone [72,73]. *Chlamydophila* co-occurred only with an unclassified *Christensenellaceae* member and did not integrate into the broader anaerobic guild; this topological isolation is consistent with the obligately intracellular lifestyle of the *Chlamydiaceae*, whose extensive genome reduction and reliance on host-derived metabolites minimises opportunities for extracellular metabolic exchange [74–77].

Functional inference by PICRUSt2, although unable to detect a significant shift in overall pathway composition for either KEGG or MetaCyc, identified three MetaCyc pathways enriched in the presence of *Chlamydiota* that together encompass the lipid-rich cell-wall and redox-buffering biology characteristic of *Actinobacteriota* [78–80]. Two of these relate directly to cell-envelope construction: mycolyl-arabinogalactan-peptidoglycan complex biosynthesis assembles the covalently linked, mycolic-acid-rich cell-wall core of these bacteria, while mono-trans, poly-cis decaprenyl phosphate biosynthesis supplies the polyprenyl lipid carrier required for assembly of the arabinan and peptidoglycan components of that same complex, so that the two pathways report on complementary steps of a single envelope-assembly programme [81–83]. The third pathway, mycothiol biosynthesis, might be more directly relevant in the context of chlamydial presence: mycothiol is the principal low-molecular-weight thiol of actinomycetes, functions as the actinobacterial analogue of glutathione, and protects against oxidative and electrophilic stress, and its increased inferred abundance in the presence of *Chlamydiota* is consistent with a pro-oxidant mucosal milieu of the kind reported in human epithelial models of chlamydial infection driven in part by IFNγ-dependent responses [17,28,79]. The convergence of mycolic-acid-rich envelope architecture and mycothiol-based redox buffering within the same expanded actinobacterial cohort therefore may suggest a coherent stress-tolerance programme selected for under the mucosal conditions accompanying chlamydial presence. These inferences are PICRUSt2 predictions from 16S rRNA data and should be interpreted with caution. Integrating the compositional, network and functional findings, our data support the view that snake cloacal *Chlamydiota* cannot be classified strictly as either pathogens or non-pathogens.

*C. serpentis* is a well-described pathogen in vipers [5,11]; *Candidatus* C. sanziniae is a recognised reptile-adapted chlamydial species that shares a phylogenetic clade with *C. pneumoniae* [6]; whilst the *Parachlamydiaceae* and *Rhabdochlamydiaceae* are CLOs for which free-living amoebae and arthropods are natural hosts and reservoirs [11,84]. The detection of *C. muridarum*, a pathogen of rodents, within a snake cloaca is most plausibly explained by recent dietary exposure to an infected rodent prey item [7,9] and raises the possibility that prey items act as a route of entry for both commensal and pathogenic *Chlamydiota* into the cloacal ecosystem. In this light, the microbiota-level changes we describe, increased dispersion, a more modular and cooperatively wired anaerobic network, and shifts in actinobacterial cell-wall and indole-related metabolism, are best interpreted as the ecological footprint of a mixed chlamydial assemblage rather than of any single organism. This view is congruent with current mammalian paradigms, in which *Chlamydia* may persist asymptomatically in the gastrointestinal tract whilst nonetheless disturbing epithelial-immune crosstalk, goblet cell biology and microbial community structure [12–14].

Our findings come with several limitations. First, the study design prevents causal inference; we cannot determine whether the presence of *Chlamydiota* drove the observed community changes or whether a permissive cloacal microbiota facilitated this presence, as proposed for human cervicovaginal risk factors for *C. trachomatis* acquisition [57]. Second, the significant association between husbandry facility and *Chlamydiota* status precluded stratified permutation schemes and introduces residual confounding that neither marginal PERMANOVA nor CAP can fully eliminate. Third, the highly unequal distribution of *Chlamydiota*-positive and -negative samples in the *Henophidia* subset restricted this study to *Caenophidia*, and our conclusions may not generalise to snake taxonomic clades with differing microbiota baselines. Finally, PICRUSt2 predictions are inferential and should be regarded as hypothesis-generating pending shotgun metagenomic and metabolomic validation.

To our knowledge, this is the first study to compare the cloacal microbiota of snakes in the presence and absence of *Chlamydiota*. The chlamydial taxa detected formed a heterogeneous assemblage spanning established reptile pathogens, CLOs, and likely dietary introductions. Although global community composition did not differ significantly between *Chlamydiota*-positive and negative caenophidian snakes, in the presence of *Chlamydiota* we observed greater within-group heterogeneity, a denser and predominantly cooperative co-occurrence network, depletion of butyrate-producing *Lachnospiraceae*, and selective expansion of an *Actinobacteriota* cohort with redox-buffering and cell-wall stress-tolerance traits. Together, these patterns sketch a revised ecological model of chlamydial presence in the snake cloaca that draws selectively on, but does not fully match, either the mammalian gastrointestinal commensalism [12–14,53,85] or the cervicovaginal dysbiosis-driven acquisition paradigms [27,28,55–57,66,67]. Longitudinal sampling to address causality, multi-omics validation of the actinobacterial stress-response hypotheses, and extension to *Henophidia* and free-ranging populations are needed to clarify the generalisability of these patterns and their implications for captive snake surveillance.

## Ethics approval and consent to participate

The collection and molecular analysis of the snake samples was approved and performed in accordance with the relevant guidelines and regulations of the Veterinary Office of Canton Zurich (authorization no. ZH010/15).

## Consent for publication

All authors consent to the publication of this manuscript.

## Author contribution

E.G.: drafting the manuscript and data analysis; S.N.: data curation and methodology; T.P.: data acquisition; S.R.: data acquisition; S.A.: data acquisition; C.B.: design of the work; N.B.: design of the work and funding acquisition; G.G.: design of the work and funding acquisition; All authors reviewed the manuscript.

## Acknowledgments

We gratefully acknowledge the sequencing facility of the DMLP at Lausanne University Hospital for their technical support and 16S rRNA amplicon sequencing, and Ms. Rizlène Dira for her assistance with sample preparation.

Category Original article Data availability

Sequencing data in the form of fastq.gz files used in this study can be accessed from the European Nucleotide Archive project accession PRJEB112904.

