## Supplementary material for "Microbial community profiles of the snake cloaca in the presence and absence of *Chlamydiota*": Fig. S1

**Category**

Original article

**Data availability**

Sequencing data in the form of fastq.gz files used in this study can be accessed from the European Nucleotide Archive project accession PRJEB112904.

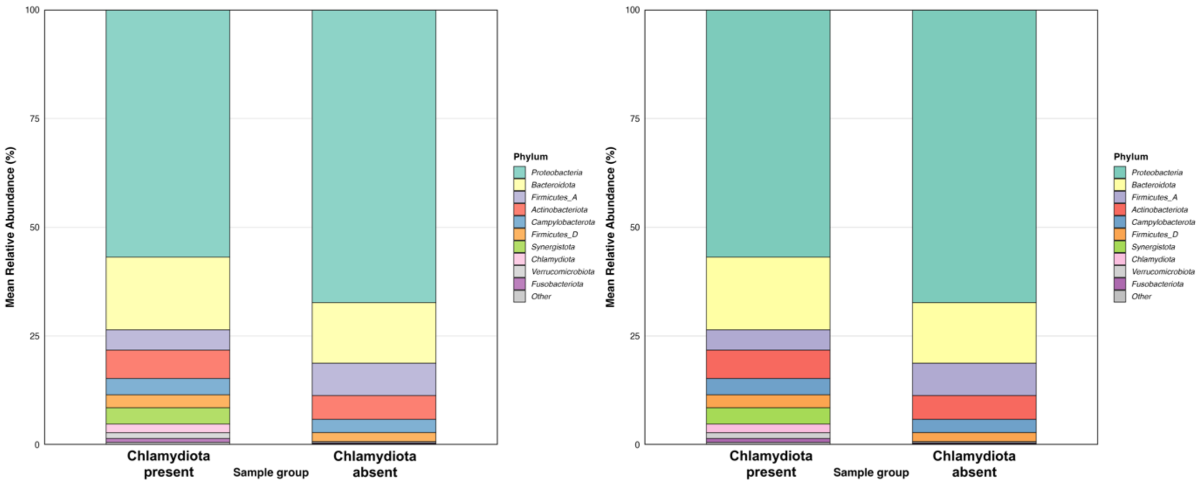


**Fig. S1. Mean relative abundance of cloacal microbiota in *Chlamydiota*-positive (n = 25) and negative (n = 27) snake samples at (a) phylum level (top 10 phyla) and (b) genus level (top 25 genera).** Stacked bars represent the mean relative abundance across samples within each group, with taxa ranked by cross-group mean abundance. Remaining low-abundance taxa are collapsed into the "Other" category. Taxa lacking genus-level annotation were assigned the name of the next higher classified rank (family or order) where available.
